# Effects of Electric Field Direction on TMS-based Motor Cortex Mapping

**DOI:** 10.1101/2024.12.10.627753

**Authors:** Ying Jing, Ole Numssen, Gesa Hartwigsen, Thomas R. Knösche, Konstantin Weise

## Abstract

**Background:** Transcranial magnetic stimulation (TMS) modulates brain activity by inducing electric fields (E-fields) that can elicit action potentials in cortical neurons. Neuronal responses to TMS depend not only on the magnitude of the induced E-field but also on various physiological factors. In this study, we incorporated a novel average response model that efficiently estimates the firing threshold of neurons based on their orientation relative to the applied E-field, thereby advancing TMS mapping for motor function.

**Methods:** We conducted a regression-based TMS mapping experiment with fourteen subjects to localize cortical origins of motor evoked potential (MEP) on the first dorsal interosseous (FDI) muscle. Firing thresholds were estimated for excitatory neurons in cortical layers 2/3 and 5 via an average response model. Regression was performed between MEPs and three E-field quantities: the magnitude (magnitude model), the normal component (cosine model), and the effective E-field, which scales the E-field magnitude based on the firing thresholds specific to the neuronal orientation (neuron model). To validate, we applied TMS to ten subjects with optimized coil placements based on these three models to determine which model could yield the highest MEPs.

**Results:** The magnitude and neuron models performed similarly, while the cosine model showed lower explained variance in regression results, required more TMS trials for stable mapping, and yielded the lowest MEP in the validation.

**Conclusion:** This study is the first to advance TMS modeling by incorporating neuron-specific factors at the individual level. Results show that on the motor cortex, the magnitude model is–as expected–a good approximation of cortical TMS effects as it shows similar results as the neuron model. In contrast, the classic cosine model exhibited lower performance and required more TMS trials for stable results, and is not recommended for future studies.

## 1. Introduction

Since the first demonstration in 1985 (Barker et al., 1985), transcranial magnetic stimulation (TMS) has become a widely used technique for non-invasively modulating brain activity in vivo. By transiently altering brain function, TMS enables causal mapping of brain structure-function relationships, using measures such as motor-evoked potentials (MEPs) to identify cortical sites responsible for specific functions (Van De Ruit et al., 2015; Gunduz et al., 2020). This capability has been widely applied in pre-surgical target selection and functional assessment following neurological injuries (Seynaeve et al., 2019; Sondergaard et al., 2021).

While TMS is FDA-cleared for cortical mapping (Cohen et al., 2022) and the treatment of several psychiatric disorders (McNamara et al., 2001; George et al., 2009), it is still largely unknown which neuron structures are primarily excited by a TMS pulse and how this localized activation leads to behavioral or psychological outcomes (Romero et al., 2019). The wide spatial distribution of TMS-induced electric fields (E-field) across large cortical areas further complicates the identification of the effective target (Bijsterbosch et al., 2012). Consequently, TMS studies often show small effect sizes (Beynel et al., 2019) and exhibit considerable variability both between and within subjects (Caulfield and Brown, 2022; Numssen et al., 2023), limiting the general efficacy of TMS in both basic research and clinical applications (Hartwigsen and Silvanto, 2023). This observed variability in TMS effects is attributed to a complex interplay of various factors, including physiological parameters across macro-, meso-, and microlevels (e.g. gyrification pattern, spatial organization of cortical neurons, neuron types, respectively), methodological parameters (e.g. pulse shape, coil geometry, stimulator intensity), and cognitive factors (e.g. response strategies, cognitive brain state) (Numssen et al., 2024; Williams et al., 2024).

To precisely map TMS effects at the macroscopic level, researchers have turned to numerical simulation of the induced E-field based on individual head and brain anatomies. Understanding the stimulation processes requires elucidating how the TMS-induced E-field modulates neuronal activity in the cerebral cortex (Laakso et al., 2018; Hartwigsen, 2015). Early mapping studies assume functionally relevant regions align with high E-field strengths (Thielscher and Kammer, 2002; Opitz et al., 2013; Aonuma et al., 2018). However, because TMS-induced E-fields affect a wide range of neural structures, the location of the highest E-field strength does not necessarily match the location of the neurons driving the observed effects (Weise et al., 2020). Later methods attempted to establish a linear relationship between E-field strength and MEP amplitude (Matthaeus et al., 2008). However, cortical response to stimulation is inherently non-linear: MEPs appear only after surpassing a threshold and plateau as E-field strength continues to increase. To address these limitations, we recently proposed a non-linear, regression-based TMS mapping method that revealed a strong, sigmoid-like relationship between local E-field magnitude and MEPs at the cortical muscle representation (Fig. 1), which is primarily located on the crowns and rims of the precentral gyrus (Weise et al., 2020; Numssen et al., 2021; Weise et al., 2023a).

**Figure 1.**
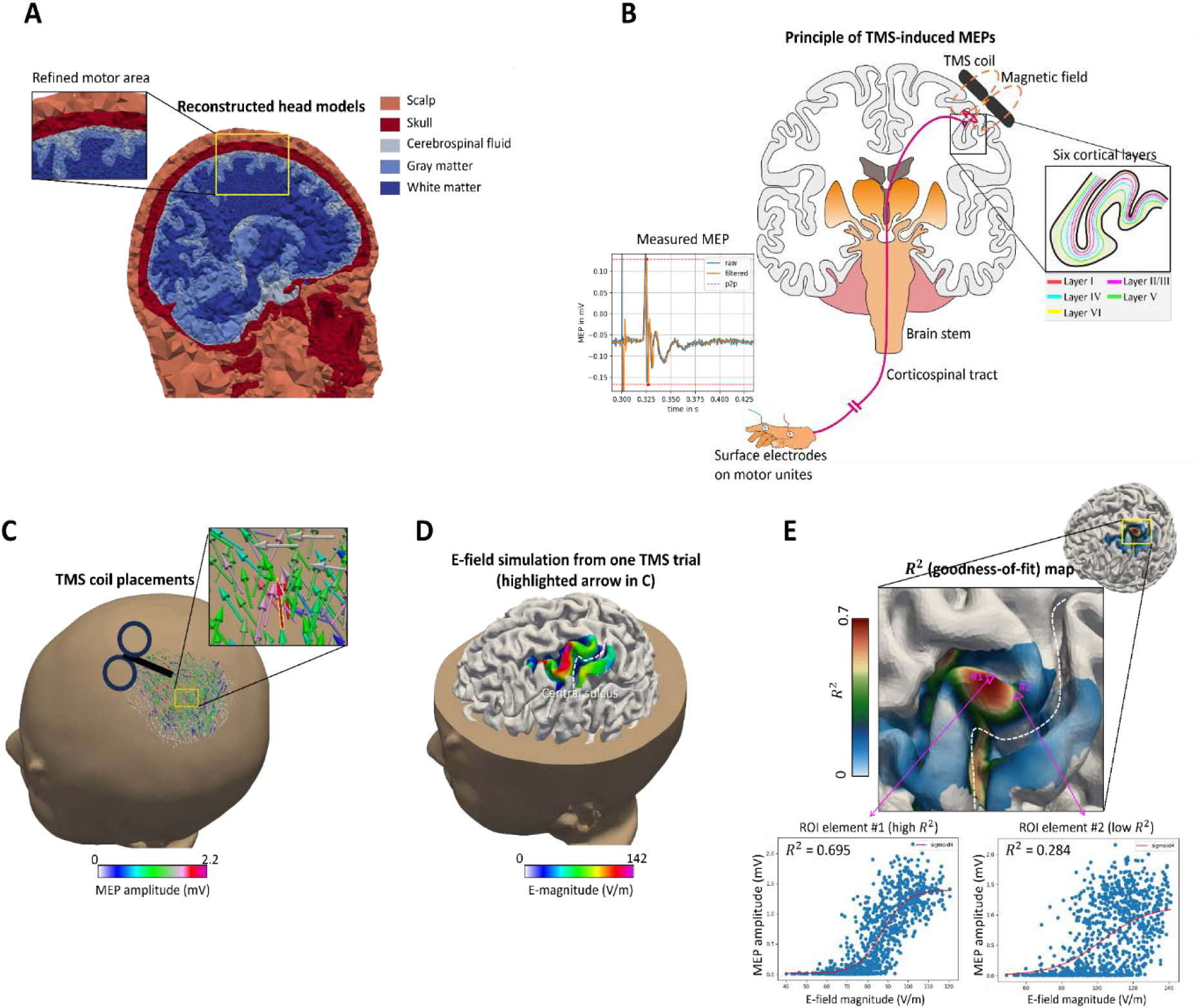
Overview of the regression-based localization approach. (A) Reconstructed head model from an example subject, consisting of five main head tissues (scalp, skull, CSF, GM, and WM). This head model is used for numerical simulations of induced E-fields. (B) Principle of TMS-induced motor evoked potentials (MEPs). The TMS coil above the M1 generates a brief current in M1 via electromagnetic induction, with induced action potentials descending along the corticospinal tract and the peripheral nerve to activate muscle units, resulting in an MEP. (C) TMS was performed on M1 with hundreds of TMS pulses from random coil positions and orientations. The color shows the peak-to-peak MEP amplitude. (D) Simulated E-fields are computed for each TMS pulse using the subject-specific head model. The color map represents the magnitude of the induced E-field from one TMS pulse (highlighted in yellow in C). (E) For each cortical element in the region of interest, different E-field quantities (|𝐸|, 𝐸_𝑒𝑓𝑓_, and |𝐸_⊥_|) were regressed against the MEP amplitudes to test different neuronal response models. This yields goodness of fit (𝑅^2^) maps that identify the most probable origin of MEPs. Only the 𝑅^2^ map from |𝐸| is shown here.

Critically, the response of neurons to TMS is not solely influenced by the magnitude of induced E-field, but also by complex interactions between E-fields and neuron morphology, as well as their spatial organization within cortical layers (Reijonen et al., 2020; Hannah and Rothwell, 2017; Di Lazzaro and Rothwell, 2014). The direction of the induced current is particularly important, as axons are activated by differences in potential along their length. Early experimental studies revealed that single neurons are more responsive to current oriented longitudinally rather than transversely across it (Rushton, 1927; Ranck Jr, 1975). This led to the well-known cortical column cosine model (Fox et al., 2004), which assumes that the depolarization threshold of axons is inversely proportional to the cosine of the angle between the external E-field and the principal axis of the neuron (Fig. 2A, B). However, this model oversimplifies neuronal activation by assuming that axonal orientation is primarily aligned with the neuron’s principal axis, typically oriented perpendicular to the cortical surface. Recent studies utilizing realistic simulations of neuronal geometry revealed that axons often exhibit complex branching, contributing to a nuanced directional sensitivity (Aberra et al., 2020; Aberra et al., 2023; Weise et al., 2023b). These simulations further found that in the motor cortex, neurons in layer 2/3 (L2/3) and layer 5 (L5) have the lowest activation thresholds, while those in layer 1 and layer 6 show almost no direct activation under typical E-field strengths (Aberra et al., 2020). While these detailed, multi-scale simulations of neuron-level responses are computationally intensive and impractical for routine TMS applications.

**Figure 2.**
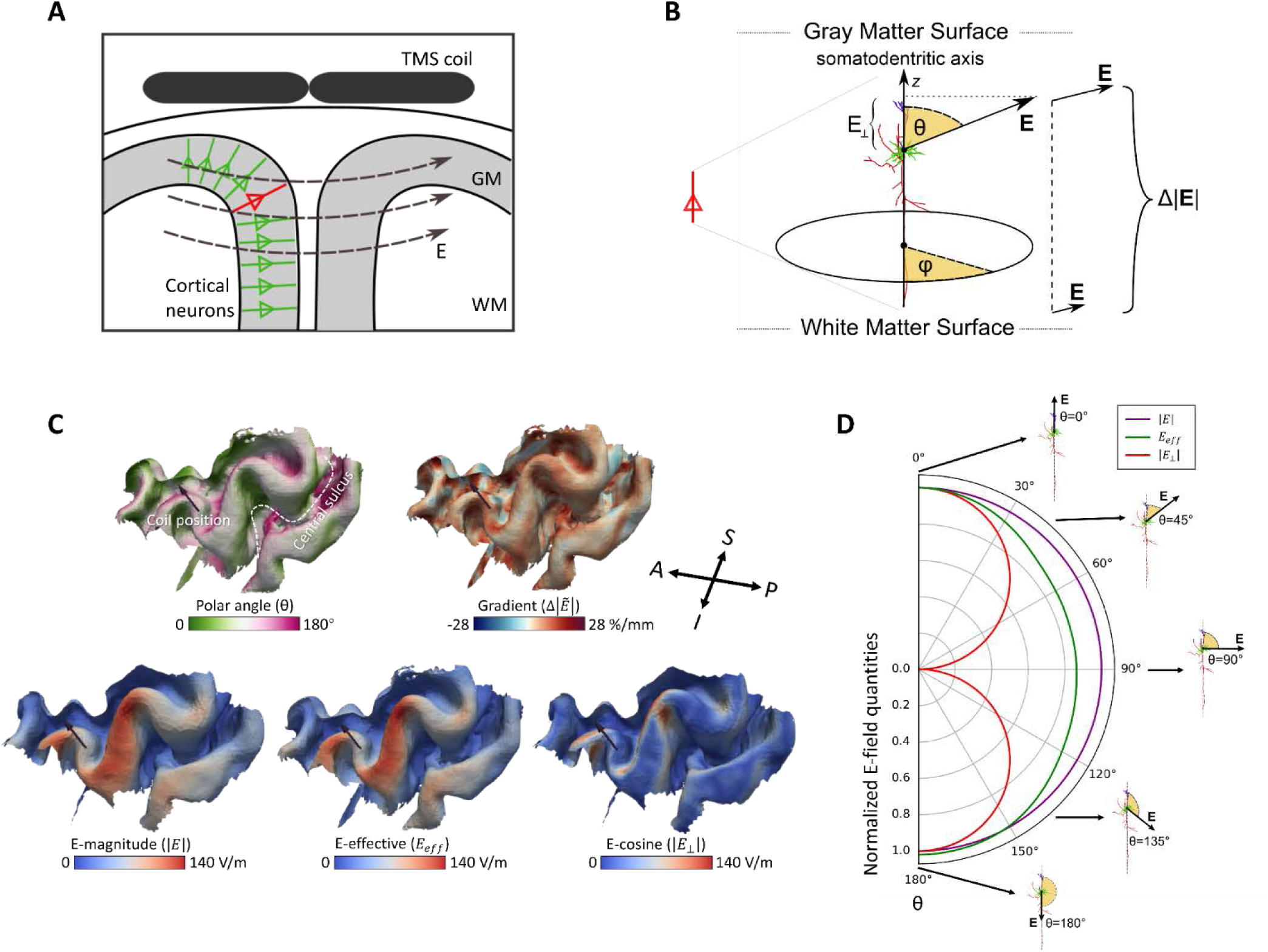
The three neuronal response models are based on different E-field quantities. (A) TMS-induced E-fields are oriented perpendicular to cortical neurons on the gyral crown but can run parallel to sulcal neurons. (B) An example neuron exposed to the E-field. θ: the angle between the E-field and the somato-dendritic axis; φ: the E- field direction in the horizontal plane perpendicular to the somato-dendritic axis. Δ|𝐸|: The gradient of the E-field magnitude between gray and white matter surfaces. |𝐸_⊥_|: The magnitude of the E-field along the somato-dendritic axis. The cosine model assumes that neurons are only responsive to |𝐸_⊥_|. (C): Allocation of polar angle θ and gradient 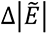 on L5, E-field magnitude |𝐸| on gray matter midlayer, effective E-field 𝐸_𝑒𝑓𝑓_ on L5, and |𝐸_⊥_| on gray matter midlayer in one example TMS pulse. The tail of the arrow represents the coil position, and its direction represents coil orientation. (D): The different neuronal response models (|𝐸|, 𝐸𝑒𝑓𝑓, and |𝐸_⊥_|) yield different neuronal stimulation (radial axis) across θ from 0° to 180° at 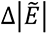 = 0. A, anterior; P, posterior; S, superior; I, inferior.

To address this, a computationally efficient model was developed that determines the averaged firing thresholds of neurons from different cortical layers by subjecting various types of cortical neurons to TMS-induced E-fields with different angles, intensities, pulse waveforms, and field gradients along the somato-dendritic axis (Weise et al., 2023b). The major advantage of this model is that it allows for a straightforward transformation from macroscopic E-field topographies to neuron firing thresholds without sacrificing modeling accuracy. The firing threshold of any given cortical location at any layer can be determined using precomputed look-up tables and interpolators, which account for the estimated polar angle θ between the E-fields and axons, as well as the field gradient 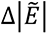 at a particular cortical location (Fig. 2B). This firing threshold quantifies the strength of the E-field required to elicit an action potential. By scaling the raw E-field magnitude |𝐸| with the firing threshold, the effective E-field, which accounts for neuronal response characteristics, can be estimated to provide a more precise estimation of TMS effects.

In this study, we incorporated this novel neuronal response model into our regression-based TMS mapping method to improve the precision of cortical muscle representation mapping. This mapping method, referred to as the neuron-enhanced model (or simply the ***neuron model***), was evaluated in comparison to two previous models: (1) the classical ***magnitude model***, as used by Numssen et al. (2021), which considers the magnitude of the E-field |𝐸| at the midlayer of gray matter, and (2) the cortical column cosine model, (or simply the ***cosine model***), which uses the normal component of E-field relative to the cortical surface. Unlike these two simplified models, the ***neuron model*** scales the E-field’s magnitude with the local firing threshold in L2/3 and L5, as neurons in these layers primarily project to the spinal cord and, thus, are the origin of MEPs (Aberra et al., 2020; Wilson et al., 2021). These three models constitute a set of ***neuronal response models***, each simulating how neurons respond to exposed E-fields (see Fig. 2D for an illustration of these three E-field quantities). We hypothesized that, compared to the magnitude and cosine models, the neuron model would explain more variance in the TMS mapping results and thus provide more precise and faster localization of cortical muscle representations, i.e., the cortical area essential for generating MEP. Whereas the neuron model predicts only moderate directional sensitivity of all neuron types (Weise et al., 2023b), the cosine model assumes a very strong directional sensitivity by stating that all axonal segments are oriented orthogonal to the cortical surface. Accordingly, we hypothesized that the magnitude model aligns better with the neuron model than the cosine model. To test these hypotheses, we conducted a second experimental session following an initial regression-based mapping session. In this validation session, TMS was performed using optimal coil placements based on all three models to determine which model yields the highest MEPs–that is, which model optimally locates the muscle representation. Additionally, we performed an extensive convergence analysis to test which model requires the fewest TMS pulses to obtain stable mapping results.

## 2. Methods

### 2.1. Subjects

**Dataset 1** was reused from Numssen et al. (2021) and consists of fourteen healthy, right-handed participants (seven females, aged 21-38 years). For a detailed description of subject recruitment, please refer to the original paper. All participants completed a TMS localization experiment in 2020, and this dataset was analyzed for both TMS localization and convergence in the current study.

To account for potential changes in motor function over the four-year interval, direct validation of localization results from dataset 1 was not feasible. Therefore, we recruited ten new participants who completed both a TMS localization session (see Section 2.4, *TMS Localization*) and a validation session (see Section 2.5, *TMS Validation Experiment*) approximately one week apart. **Dataset 2** includes these ten healthy, right-handed participants (five females, aged 24-43 years), with a mean laterality index of 71 (SD = 15) as measured by the Edinburgh Handedness Inventory. None of these participants had contraindications to TMS, were on regular medication, or had a history of psychiatric or neurological diseases.

All participants in both datasets provided written informed consent to participate prior to the experiment. The studies were approved by the local Ethics committee of the Medical Faculty of Leipzig University.

### 2.2. Imaging

Structural and diffusion-weighted magnetic resonance imaging (MRI) data were collected using a 3 Tesla Siemens Skyra fit scanner with a 32-channel head coil. To segment the main tissues of the head for E-field calculations (Fig. 1A), T1- and T2- weighted images were acquired with following parameters: T1 MPRAGE sequence with 176 sagittal slices, matrix size = 256 × 240, voxel size = 1 × 1 × 1 mm^3^, flip angle 9°, TR/TE/TI = 2300/2.98/900 ms; T2-weighted with 192 sagittal slices, matrix size = 512 × 512, voxel size = 0.488 × 0.488 × 1 mm^3^, flip angle 120°, TR/TE = 5000/395 ms. The T1-weighted image was also used for neuronavigation during TMS. Diffusion MRI with 88 axial slices, matrix size = 128 × 128, voxel size = 1.719 × 1.719 × 1.7 mm^3^, TR/TE = 80/7000 ms, flip angle 90°, 67 diffusion directions, b-value 1000 s/mm^3^ was acquired for the estimation of the conductivity anisotropy in the gray and white matter. If existing scans in the image database were less than two years old, they were utilized instead.

### 2.3. Electromyography

Electromyograms (EMGs) from the right first dorsal interosseous (FDI) muscle were recorded using pairs of Ag-AgCl surface electrodes in a standard belly-tendon montage (Kleim et al., 2007) (Fig. 1B). The raw signal was amplified with a patient amplifier system (D-360, DigitimerLtd., UK, Welwyn Garden City), bandpass filtered from 10 Hz to 2 kHz, and recorded with an acquisition interface (Power1401 MK-II, CED Ltd., UK, Cambridge) at a 4 kHz sampling rate using Signal software (CED Ltd., version 4.11). MEPs were calculated as peak-to-peak amplitudes within a time window of 18 to 35 ms after the TMS pulse.

### 2.4. TMS Localization

#### 2.4.1. TMS Protocol

TMS pulses were applied using a MagPro X100 stimulator (MagVenture, firmware Version 7.1.1) with an MCF-B65 figure-of-eight coil. A TMS navigation system (Localite software, Germany, Sankt Augustin; camera: Polaris Spectra, NDI, Canada, Waterloo) was employed to guide motor threshold (MT) hunting and to record the coil’s orientations and positions in real-time during the localization experiment.

We manually measured the resting MT (rMT) for the FDI while participants were at rest, with earplugs provided to prevent hearing damage during TMS. The stimulation site was first determined by locating the hand knob on the precentral gyrus (Yousry et al., 1997) based on individual T1 images. The motor hotspot was then identified by systematically varying the coil’s tilt, rotation, and location until a placement was found that elicited stable MEPs. An estimate of the rMT was determined at the motor hotspot as the lowest stimulator intensity required to produce MEPs with an amplitude of at least 50 𝜇V in at least 5 out of 10 consecutive TMS pulses (Rothwell et al., 1999).

In dataset 1, we utilized the data described by Numssen et al. (2021). In brief, to localize the muscle representation of the FDI, 900–1100 single biphasic TMS pulses were applied at 150% rMT with an inter-stimulus interval (ISI) of 5 s to the left motor area. For each stimulation, coil position and orientation were randomly selected within a 2 cm radius and the angles to approximately ± 60° in relation to the optimal coil placement (Fig. 1C). The purpose of this randomization was to increase E-field variability and minimize cross-correlations between induced E-fields. All coil placements and corresponding MEPs from the FDI muscle were recorded.

Given that participants’ motor functions might have slightly changed after four years, we recruited ten new subjects for dataset 2 to perform the localization experiment again. We followed the same stimulation protocol as described above but with only 500 random TMS pulses. As shown in our convergence analysis (see section 3.2, *Convergence Analysis*), a minimum of 300 pulses was sufficient to achieve stable localization results.

#### 2.4.2. E-field Simulation

E-fields induced for each TMS pulse were computed using SimNIBS v4.1.0 (Saturnino et al., 2019; Thielscher et al., 2015) with high-resolution anisotropic finite element models (FEMs). To simulate E-fields, individual head models were first reconstructed via the CHARMpipeline, which integrates FreeSurfer outputs for a more accurate representation of smaller sulci in the head meshes (Puonti et al., 2020). A refined region of interest (ROI) covering the somatosensory cortex (BA1, BA3), primary motor cortex M1 (BA4), and dorsal premotor cortex PMd (BA6) was defined based on the fsaverage template to improve the numerical resolution around the ROI (Fig. 1A) (Numssen et al., 2021). The final head models were composed of approximately 0.95 × 10^6^ nodes and 5.49 × 10^6^ tetrahedra (average volume: approximately 0.94 mm^3^ in the cortex, and 0.05 mm^3^ in the refined cortical ROI in the motor area). Six tissue types were included with standard conductivity estimates: white matter (𝜎_𝖶𝑀_ = 0.126 S/m), gray matter (𝜎_𝐺𝑀_ = 0.275 S/m), cerebrospinal fluid (𝜎_𝐶𝑆𝐹_ = 1.654 S/m), bone (𝜎_𝐵_ = 0.01 S/m), skin (𝜎_𝑆_ = 0. 465 S/m), and eyeballs (𝜎_𝐸𝐵_ = 0.5 S/m) (Thielscher et al., 2015). WM and GM were assigned anisotropic conductivities, while the four other tissues were treated as isotropic.

Given that the magnitude and cosine models consider GM as a uniform entity and disregard the cytoarchitectonic differences across cortical layers, we first calculated E-fields only at the midlayer between GM and WM surfaces to avoid boundary effects of the E-field due to conductivity discontinuities (Fig. 1D). The E-field interpolation employs the super-convergent patch recovery method to determine the E-field at the GM center via linear interpolation (Saturnino et al., 2019).

To incorporate detailed geometric information across different cortical layers, we added cortical layers to the motor cortex ROI in each head model. These layers were defined by their normalized depths, ranging from 0 (gray matter surface) to 1 (white matter surface), based on estimates from primate motor cortex slices (Garcja-Cabezas and Barbas, 2014). The normalized depths for the centers of these layers are 0.06 for layer 1, 0.4 for layers 2/3, 0.55 for layer 4, 0.65 for layer 5, and 0.85 for layer 6 (Aberra et al., 2020; Weise et al., 2023b). We linearly interpolated the positions of these layers between the white and gray matter surfaces using the vertex positions of the two surfaces. E-fields in layer 2/3 and layer 5 were then interpolated using SimNIBS, as they were for the midlayer. The number of ROI elements was about 0.95 × 10^4^ at the midlayer, and 2.78 × 10^4^ for layer 2/3 and layer 5.

#### 2.4.3. Firing Threshold and Effective E-field Calculation

The firing threshold of neurons, defined as the minimum E-field intensity that elicits action potentials, was estimated using the average threshold model (Weise et al., 2023b). This model provides pre-computed look-up tables for neuron firing threshold across a wide range of neuronal types and spatial orientations. Specifically, compartments of layer 2/3 and layer 5 pyramidal cells were simulated based on data from the Blue Brain Project (Markram et al., 2015). The simulated neurons were then exposed to E-fields from different directions and strengths to determine the thresholds for each cell across all possible E-field configurations. The average threshold model was derived by averaging these thresholds over all compartment models and azimuthal orientations. The model’s accuracy was validated by comparing its results with computationally intensive reference simulations, which utilized a high-resolution, realistic head model containing a large number of neurons located within the motor cortex.

To quantify the firing threshold of any given elements within the cortical ROI, we first calculated two key parameters for layer 2/3 and layer 5: ⅰ) polar angle θ: the relative angle between induced E-fields and the surface normal (somato-dendritic axis) of the cortical layers, ranging from 0° to 180°, ⅱ) field decay 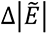 the relative change of the E-field magnitude along the somato-dendritic axis per unit length in %/mm (Fig. 2B, C). It was calculated by extracting the E-field at a normalized depth of 10% of the distance between the current layer and the GM or WM surface, respectively. Firing thresholds for each element in the ROI across layers were then determined using the average threshold model, which provides a look-up table that defines the firing threshold in relation to the polar angle θ and field decay 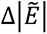.

Subsequently, we calculated the effective E-field (𝐸_𝑒𝑓𝑓_) by dividing the E-field magnitude (|𝐸|) by the normalized firing threshold (𝑡̃) for each ROI element:

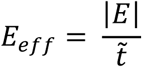

The normalized firing threshold is obtained by dividing the original firing threshold for the given polar angle and gradient by the value for θ = 0 and 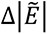 = 0. By scaling |𝐸| with the firing threshold, we quantified the effective E-field to which neurons respond proportionally to their intrinsic excitability. This method accounts for the variability of neuronal excitability across different cortical layers and brain elements, allowing for a more accurate quantification of the stimulation effects on neurons.

#### 2.4.4. Cortical Localization

We applied a regression-based localization method with random coil placements to identify muscle representations in the motor cortex where the neural populations are responsible for the TMS-induced MEPs. The rationale behind this method is that at the cortical origin of the MEPs, a clear sigmoidal relationship must exist between induced E-field and MEP amplitude. This sigmoidal or s-shaped curve, known as the input-output curve (IO) curve, captures how targeted motor neurons respond to varying stimulation strengths. At low stimulation strength (low |𝐸|), MEP amplitudes remain below a baseline or noise floor, which is likely due to unrelated neural and muscular sources. As the stimulation strength increases, MEP amplitudes rise monotonically until reaching saturation, forming the high-side plateau of the sigmoidal IO curve (Li et al., 2022). Thus, we identified the cortical origin by selecting the location with the highest goodness-of-fit (GOF) for the sigmoidal regression (Fig. 1E).

We propose that the regression based on the ***neuron model*** is the most accurate approximation of the effects of TMS-induced E-fields on MEPs. This method correlates 𝐸_𝑒𝑓𝑓_ at layer 2/3 and layer 5 pyramidal cells with the measured MEP amplitudes using a sigmoidal function. As the 𝐸_𝑒𝑓𝑓_accounts for axonal orientations and other neuronal morphologies, this method should describe the TMS-induced effects on brain activities in the most realistic manner.

Previously employed methods like the ***magnitude model*** and the ***cosine model*** can be viewed as approximations of the neuron model’s directional profile. To test these approaches against the neuron model, |𝐸| and |𝐸_⊥_| on the midlayer were also used to regress with MEPs. The magnitude model assumes that the magnitude of the E-field (|E|) best predicts the MEPs, whereas the cosine model considers the normal component of the E-field towards the cortical surface normal (𝐸_⊥_) responsible for the resulting MEPs (Fig. 2B). Here, we assume that the same magnitude of 𝐸_⊥_with opposite direction alongside the somato-dendritic axis yields the same neural activation, and, accordingly, we utilize the absolute value of 𝐸_⊥_ (|𝐸_⊥_|) in our calculations. To reduce cases where 𝐸_⊥_ becomes negative, TMS coil was positioned within a ± 60° range relative to the PA 45° (posterior-anterior 45 degrees) orientation, which aligns well with the typical axonal orientation in motor cortical neurons. We hypothesize that the magnitude model provides a better approximation compared to the cosine model based on the greater similarity of its angular profile to that of the neuron model (Fig. 2D).

A sigmoidal IO curve was used to characterize the relationship between the three kinds of E-field quantities (Fig. 2D) with the elicited MEPs for every element within the ROI, respectively:

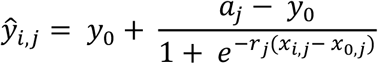

Here, 𝑥_𝑖,𝑗𝑗_ is the E-field of TMS pulse 𝑖 (1 ≤ 𝑖 ≤ 𝑁_𝑇𝑀𝑆_) at cortical element *j* (1 ≤ 𝑗𝑗 ≤ 𝑁_𝑒𝑙𝑚𝑠_), and 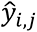is the fitted MEP, 𝑎 defines the saturation amplitude, 𝑟 is the slope, 𝑥_0_ is the location of the turning point on the 𝑥-axis, and 𝑦_0_ denotes the floor value that corresponds to measurement noise.

The site of effective stimulation can be quantified by the GOF, which is highest at the cortical site that houses the relevant neuronal populations (Fig. 1E). We assessed the element-wise GOF by the coefficient of determination 𝑅^2^:

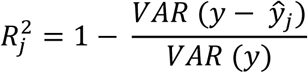

Here, 𝑦 is the measured MEP, and 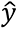 is the fitted MEP. Higher 𝑅^2^ denotes better fitting results.

We then compared the 𝑅^2^ results obtained from the 𝐸_𝑒𝑓𝑓_ at L2/3 and L5 with those derived solely from the |𝐸| and |𝐸_⊥_| by non-parametric Wilcoxon signed-rank test. Due to the spatial autocorrelation (or “smoothness”) of the E-field, neighboring deep areas may also exhibit spurious correlations between E-fields and MEP amplitudes. To account for this, we limited 𝑅^2^ peaks to regions where at least 25% of TMS trials had |𝐸| exceeding 40 V/m. Additionally, we restricted 𝑅^2^ peaks to the precentral gyrus as it houses the primary motor cortex, responsible for motor control. Limiting the analysis to this region ensures that the observed 𝑅^2^ peaks are directly relevant to motor function and TMS stimulation effects.

#### 2.4.5. Convergence Analysis

We conducted a convergence analysis on these three models using dataset 1 to evaluate their stability and efficiency by determining the minimum number of TMS trials needed for a robust cortical localization, following the method of Numssen et al. (2021). For each subject, we first randomized the sequence of all TMS trials. Sequential regressions were then computed for 𝑛 = 10 to 800 TMS pulses to quantify the pulse-by-pulse change of the resulting 𝑅^2^map. This process was repeated 100 times with random initializations to ensure that 𝑅^2^maps converged to a stable solution regardless of initial parameter values. The analysis starts at 𝑛 = 10 and ends at 𝑁 = 800 based on previous findings (Numssen et al., 2021), indicating that localization results do not converge before the 10th pulse and generally stabilize around the 200th pulse. these resulting in 79100 𝑅^2^maps (100 random sequences × 791 zaps).

These 𝑅^2^maps were then used to compute two metrics: 1) the normalized root mean square deviation (NRMSD) to measure the overall shape similarity of the cortical localization 𝑅^2^ map; and 2) the geodesic distance of 𝑅^2^peak to quantify the accuracy of hotspot identification. Convergence for both metrics was evaluated against the full set of stimulations as a proxy for the ground truth, as well as against the previous solution from 𝑛 – 1 stimulation, to measure the speed of convergence.

The NRMSD between the 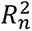 map for 𝑛 stimulations and the reference map 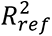 was calculated as follows:

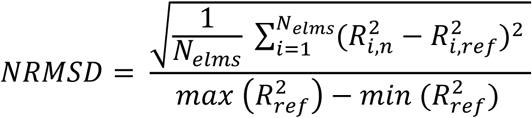

Here, 𝑖 denotes the elements (1 ≤ 𝑖 ≤ 𝑁_𝑒𝑙𝑚𝑠_) in the motor ROI, and the reference map 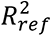 is either the 𝑅^2^ at 𝑁 stimulations or at 𝑛 − 1 stimulations.

We set convergence thresholds at 5% for NRMSD and 1 mm for the geodesic distance between hotspots. The minimum number of stimulations needed to meet these criteria against the reference solution is determined as the threshold for achieving stable cortical localizations. This convergence analysis provides both a practical stopping criterion for online TMS experiments and a method for post-hoc evaluation of the overall effectiveness of the mapping procedure’s effectiveness. To obtain robust convergence estimates for each subject, we averaged the minimum number of stimulations meeting the convergence criteria across 100 randomized sequences for each stimulation count (from 10 to 800).

To determine which model yields the most stable cortical localization (𝑅^2^map), we compared the mean minimum number of stimulations required to meet the NRMSD and geodesic distance thresholds across the three models at the group level using the Friedman test, a non-parametric repeated measures ANOVA. Conover’s Post-hoc analysis was then conducted to identify significant differences between the models, with multiple-comparison corrections for pairwise comparisons. For the neuron model, we only used data from L5, as it produced similar localization results to those from L2/3.

### 2.5. TMS Validation Experiment

After identifying the neuronal populations responsible for TMS-induced MEPs, i.e., the hotspots with the highest 𝑅^2^ based on three models in ten subjects in dataset 2, we conducted a second TMS experiment session to validate these results.

For each subject and each of the three models, we determined the optimal coil placement, i.e., coil position and orientation, that yielded the maximum 𝐸_𝑒𝑓𝑓_, |𝐸|, or |𝐸_⊥_| at the 𝑅^2^peak, respectively. This was done using an exhausting search procedure, which involved projecting the hotspots onto the head surface and defining search grids centered on these projected sites with a radius of 20 mm, angle range of 360°, spatial resolution of 2 mm, and angle resolution of 4°, resulting in 27450 coil configurations for each model (Fig. 6A). E-field distributions were then simulated within the motor ROI for each coil configuration. The coil placements that yield the maximum 𝐸_𝑒𝑓𝑓_from L5, |𝐸|, and |𝐸_⊥_| at the corresponding 𝑅^2^ hotspots were imported to the neuronavigation system (Kalloch and Numssen, 2022). This optimization procedure is implemented in SimNIBS (https://simnibs.github.io/simnibs) and has been previously applied by Weise et al. (2020) and Numssen et al. (2021).

Additionally, we included four surrounding coil placements that shared the same optimal coil orientation as the magnitude-model-based coil orientation but were shifted 10 mm in the superior, posterior, inferior, or anterior directions relative to the original magnitude coil placement. These four coil placements serve as control conditions (Fig. 6B).

To validate the mapping results, we applied TMS at each of the seven coil placements using the same protocol as outlined in the localization experiment (see Section 2.4.1, *TMS Protocol*), with single biphasic pulses at a 5-second ISI. The stimulation intensity was set to 120% rMT instead of 150%, as this level typically yields robust MEPs while remaining highly sensitive to minor intensity changes (Houdayer et al., 2008; Souza et al., 2022). Fifty TMS trials were collected for each target to obtain a stable MEP estimate from the FDI muscle (Goldsworthy et al., 2016). The order of coil placements was randomized per subject to mitigate sequence effects. The averaged MEP peak-to-peak amplitudes were then compared across these seven coil placements by the non-parametric Friedman test. We expected the magnitude and neuron models to yield higher MEP amplitudes than other placements.

## 3. Results

### 3.1. Localization Maps

We performed a sigmoidal regression between the three E-field quantities and TMS-elicited MEPs to localize the cortical sites responsible for the observed MEP in the right FDI muscle. The effective stimulation site is assumed to have the highest 𝑅^2^ value at the cortical location housing the relevant neuronal populations. Subject-wise localization results for the magnitude, neuron, and cosine models are presented in Fig. 3 and Table S1.

**Figure 3.**
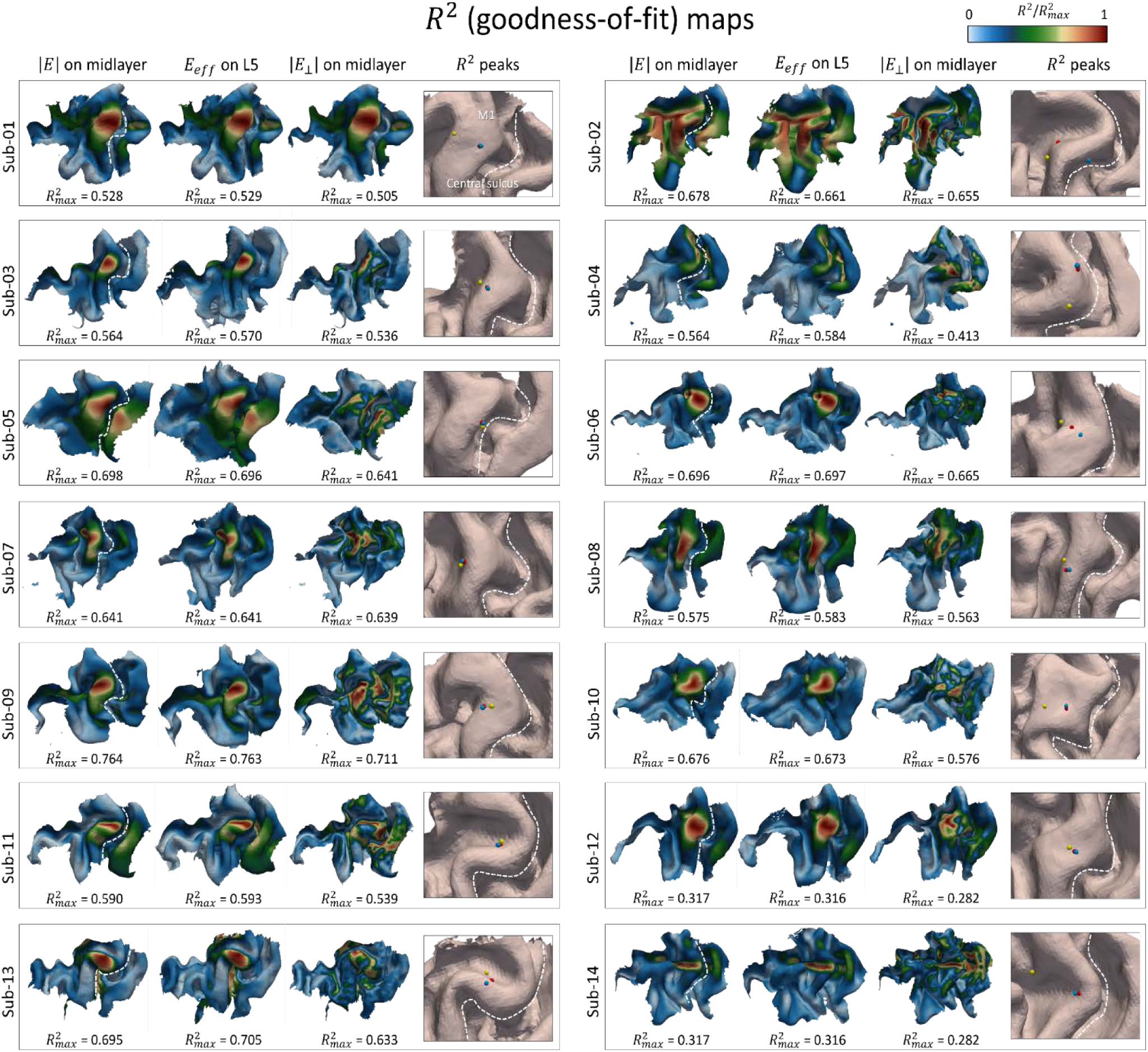
Individual motor mapping results for magnitude, neuron, and cosine response models. First three columns: normalized 𝑅^2^ maps from magnitude model (|𝐸|) on gray matter midlayer, neuron model (𝐸_𝑒𝑓𝑓_) in L5, and cosine model (|𝐸⊥|) on midlayer. Maximum 𝑅^2^ values range from 0.28 to 0.77. The cosine model yields lower 𝑅^2^values compared to the magnitude and neuron model, thus showing poorer performance in localizing cortical muscle representations. The last column highlights the identified hotspot for all three models. The locations derived from the magnitude and neuron models are similar, while the hotspot from the cosine model is significantly distant.

The overall shape of the normalized 𝑅^2^ maps is very similar between the magnitude and neuron models, which incorporated neuron morphologies from L2/3 and L5. Their 𝑅^2^ peak values are also not significantly different (mean_mag_ = 0.598; mean_neuron_L2/3_ = 0.600; mean_neuron_L5_ = 0.600; magnitude vs. neuron (L2/3): *Z* = 43.5, *p* = 0.58; magnitude vs. neuron (L5): *Z* = 49.5, *p* = 0.90, Wilcoxon signed-rank test). Given the similarity in results between L2/3 and L5, we focused on the L5 data for subsequent analyses.

In contrast, the cosine model shows significantly lower 𝑅^2^ peak values compared to the magnitude model (mean_cos_ = 0.551; *Z* = 0.0, *p* = 1.2 × 10^-4^). A test statistic of *Z* = 0.0 indicates that for all data pairs, the 𝑅^2^ peak values from the cosine model are consistently lower than those from the magnitude model. Similarly, the comparison between the cosine and neuron models also shows a significant difference (*Z* = 0.0, *p* = 1.2 × 10^-4^), further confirming that the cosine model provides a less accurate functional localization of the cortical origin of the elicited MEPs. Moreover, the 𝑅^2^ maps from the cosine model appear noisier than those from the magnitude and neuron models, indicating reduced spatial coherence and accuracy in cortical localization. Fig. 4A provides both individual and group mean 𝑅^2^peak values for all models across all subjects.

**Figure 4.**
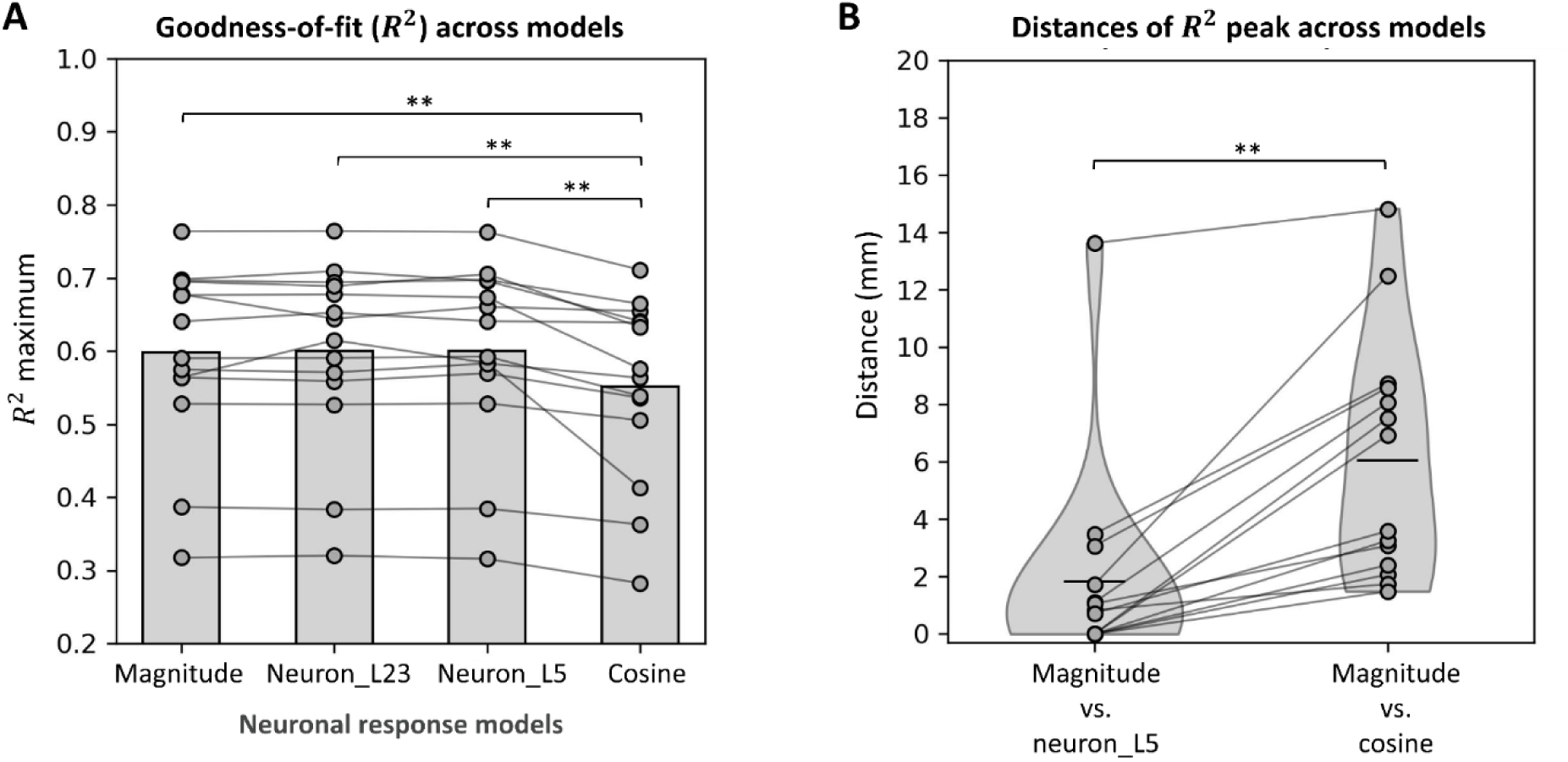
Comparisons of 𝑅^2^ peak values and locations across neuronal response models. (A) The cosine model exhibits the lowest 𝑅^2^ peak values, indicating that the normal component of E-fields explains less variance of MEPs compared to other models. (B) Hotspot locations identified by the magnitude model are significantly closer to those identified by the neuron model than with the cosine model. Black lines represent mean distances between models. \**p* < 0.05; \*\**p* < 0.01.

Likewise, the locations of maximum 𝑅^2^ values are significantly closer between the magnitude and neuron model compared to the distance between the magnitude and cosine models (Fig. 4B, Table S2, *Z* = 0.0, *p* = 1.2 × 10^-4^). This highlights the strong agreement between the magnitude and neuron models in identifying cortical hotspots, while the cosine model shows more divergence.

### 3.2. Convergence Analysis

We conducted a convergence analysis to determine which model requires fewer TMS trials to achieve a stable and correct cortical localization, quantified as the overall similarity of 𝑅^2^maps (NRMSD) and geodesic distance of 𝑅^2^peak locations (Fig. 5). Both metrics were used to compare each model’s n-th solution to both the ground truth (N stimulations) and the previous solution (n-1). Overall, both the magnitude and neuron models converged faster than the cosine model.

**Figure 5.**
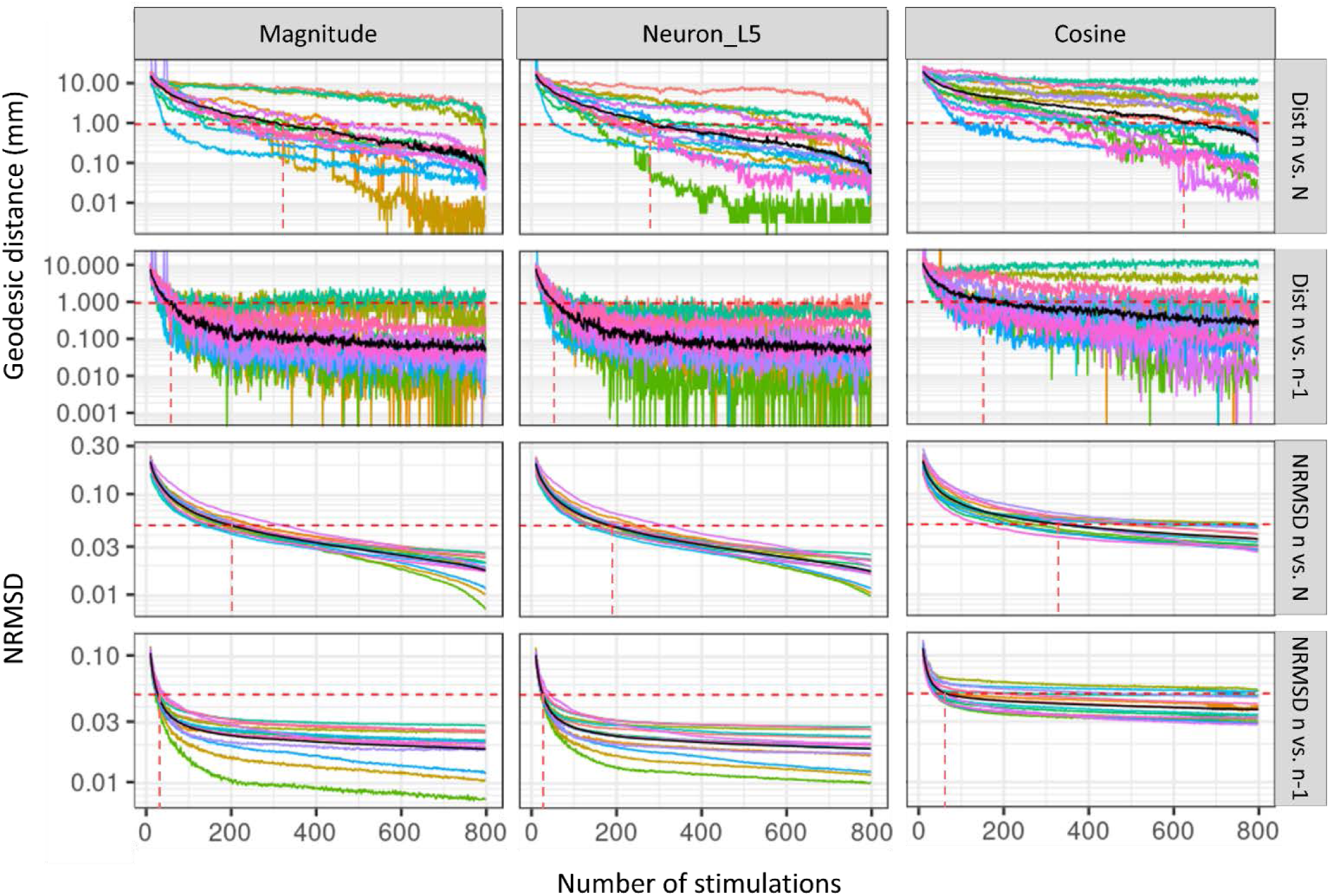
Comparison of motor localization (𝑅^2^ map) convergence across models. The magnitude and neuron models converge faster than the cosine model, i.e., they require fewer TMS pulses to achieve stable mapping results. The first two rows show the geodesic distance of the identified hotspot from the current n stimulations compared to both the 9reference solution (N) and the previous solution (n-1 stimulations). The last two rows show map similarity, represented by NRMSD, between the maps from n stimulations and both the reference solution (N) and the previous solution (n-1). Colored lines: subject-wise average convergence across 100 randomizations. Black lines: the grand mean. Red dashed line: the number of stimulations required to reach a 1 mm geodesic distance and 5% NRMSD relative to the reference solution N and previous solution n-1, respectively. Convergence plots were cut at 800 pulses.

For the geodesic distance between 𝑅^2^ peak locations from the n-th solution to the ground truth, the magnitude model required 309 TMS trials, the neuron model required 282 stimulations, and the cosine model required 622 trials to reach the 1 mm threshold. A Friedman test revealed a marginal difference among these three models in terms of the minimum number of stimulations needed for stable 𝑅^2^ peak locations (𝜒^2^ = 5.57, *p* = 0.06). Conover’s post hoc test, corrected for Bonferroni multiple comparisons, further indicated that the cosine model required significantly more stimulations to reach stability compared to both the magnitude model (*Z* = 2.25, *p* = 0.045) and neuron models (*Z* = 2.57, *p* = 0.02). No significant difference was found between the magnitude and neuron models (*Z* = -0.32, *p* = 1.00).

When comparing the current solution n with the previous n-1 solution, the magnitude model required 59 trials, the neuron model 57 trials, and the cosine model 152 trials to achieve stable peak distances. A Friedman test showed a significant difference across models (𝜒^2^ = 18.58, *p* < 0.0001). Post hoc analysis confirmed that the cosine model needed significantly more trials to reach continuous localization results than both the magnitude (*Z* = 4.79, *p* < 0.0001) and neuron models (*Z* = 4.89, *p* < 0.0001). No significant difference was found between the magnitude and neuron models (*Z* = -0.10, *p* = 1.00).

In terms of NRMSD, which quantifies the overall similarity of two 𝑅^2^ maps, the magnitude and neuron models required 200 and 191 trials, respectively, to reach the 5% deviation threshold compared to the full set of N stimulations. The cosine model, however, required 278 trials to reach the same threshold. A Friedman test revealed a significant difference across models (𝜒^2^ = 24.14, *p* < 0.0001), and post hoc analysis confirmed that the cosine model converged significantly slower than both the magnitude (*Z* = 3.92, *p* < 0.0001) and neuron models (*Z* = 4.83, *p* = 0.0005). Additionally, 31 and 28 trials were needed for the magnitude and neuron models to produce a similar localization pattern compared to the previous solutions, while the cosine model required 61 trials. A Friedman test showed a significant difference across models (𝜒^2^ = 25.08, *p* < 0.0001), with post hoc analysis indicating that the cosine model again converged significantly slower than both the magnitude (*Z* = 5.75, *p* < 0.0001) and neuron models (*Z* = 6.82, *p* < 0.0001).

These findings highlight that both the magnitude and neuron models converge more quickly and require fewer TMS trials to achieve stable and continuous localization patterns, while the cosine model lags behind, requiring significantly more trials to reach the same level of stability. This suggests that the magnitude and neuron models provide more efficient and reliable cortical localization compared to the cosine model.

Comparison of motor localization (𝑅^2^ map) convergence across models. The magnitude and neuron models e faster than the cosine model, i.e., they require fewer TMS pulses to achieve stable mapping results. The rows show the geodesic distance of the identified hotspot from the current n stimulations compared to both rence solution (N) and the previous solution (n-1 stimulations). The last two rows show map similarity, ted by NRMSD, between the maps from n stimulations and both the reference solution (N) and the previous (n-1). Colored lines: subject-wise average convergence across 100 randomizations. Black lines: the grand ed dashed line: the number of stimulations required to reach a 1 mm geodesic distance and 5% NRMSD to the reference solution N and previous solution n-1, respectively. Convergence plots were cut at 800

### 3.3. Validation

To validate the cortical mapping results from the three neuronal response models, we calculated the optimal coil placements on **Dataset 2** for the identified cortical hotspots. Experiments were then conducted with the guidance of these calculated optimal coil placements, and MEP amplitudes were measured to evaluate the accuracy and reliability of each model’s mapping. In addition to the three optimized coil positions derived from the magnitude, neuron, and cosine models, four adjacent coil placements were tested as control conditions to provide comparative data (Fig. 6C, Table S5).

**Figure 6.**
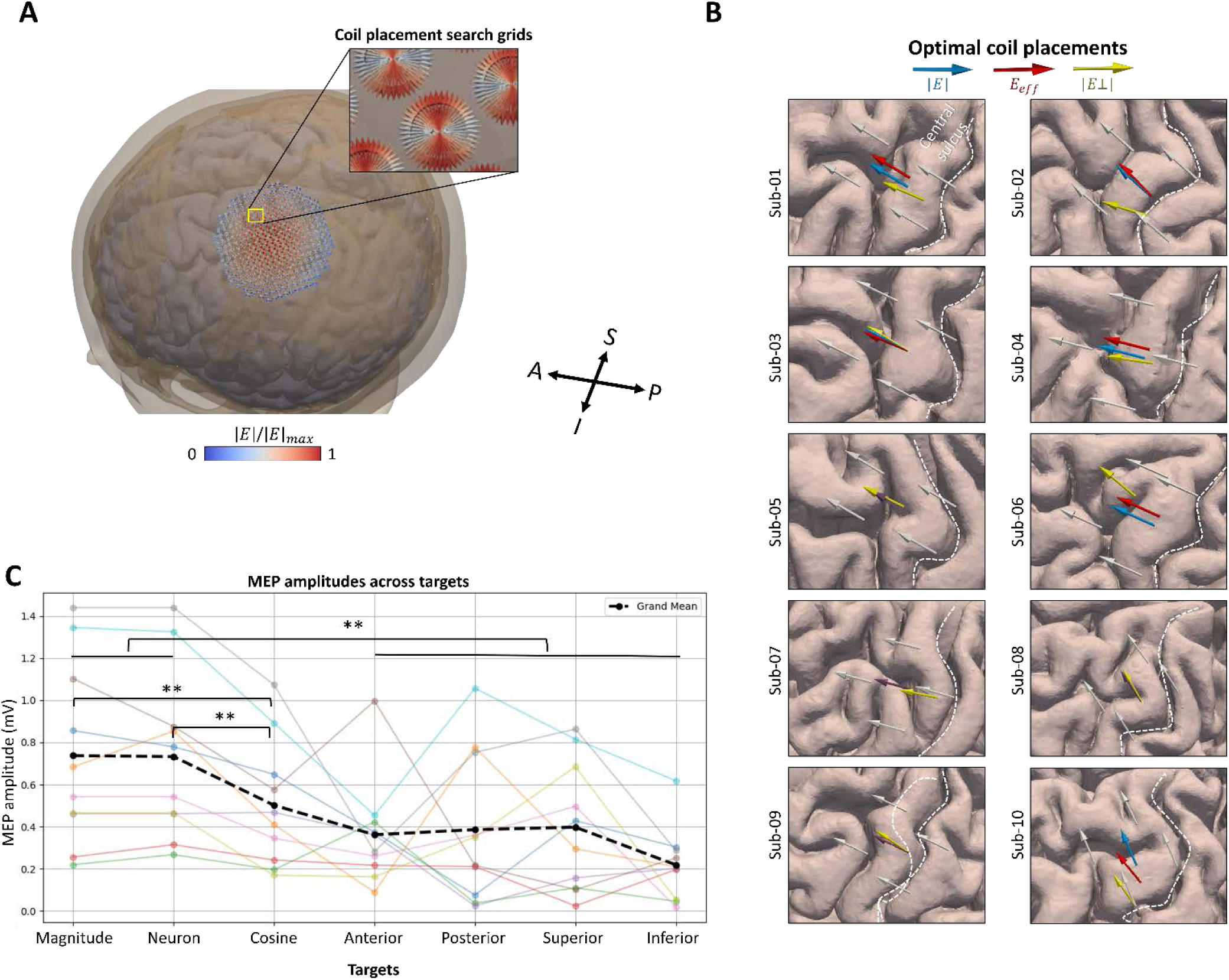
Experimental validation of the identified muscle representations across the three activation models. (A) An exhaustive search procedure was performed, testing 27450 coil placements (arrows) for each model to find the coil placement that maximizes the E-field quantity at the corresponding 𝑅^2^peak hotpot. (B) Optimal coil placements for the three models, along with four surrounding control sites at the individual level. The tail of each arrow indicates the coil location, while the arrow’s direction represents coil orientation. Purple arrow: the overlap between coil placements from magnitude (blue) and neuron (red) models at L5. (C) On average (black dashed line), the magnitude and neuron models yielded higher MEP peak-to-peak amplitudes compared to the cosine model and the four surrounding control sites, implying better localization of the cortical muscle representations. Colored points: subject-wise mean MEP amplitudes for 50 TMS pulses. Black dashed lines: the grand mean. A, anterior; P, posterior; S, superior; I, inferior. \**p* < 0.05; \*\**p* < 0.01.

Experimental validation of the identified muscle representations across the three activation models. (A) An ve search procedure was performed, testing 27450 coil placements (arrows) for each model to find the coil nt that maximizes the E-field quantity at the corresponding 𝑅^2^peak hotpot. (B) Optimal coil placements for models, along with four surrounding control sites at the individual level. The tail of each arrow indicates location, while the arrow’s direction represents coil orientation. Purple arrow: the overlap between coil nts from magnitude (blue) and neuron (red) models at L5. (C) On average (black dashed line), the magnitude ron models yielded higher MEP peak-to-peak amplitudes compared to the cosine model and the four ing control sites, implying better localization of the cortical muscle representations. Colored points: subject- an MEP amplitudes for 50 TMS pulses. Black dashed lines: the grand mean. A, anterior; P, posterior; S, superior;.

We first conducted the Shapiro-Wilk test to assess the distribution of MEPs across the seven targets. Results indicated that six of the seven targets met the normality assumption, with only the anterior target deviating slightly from normality. Therefore, we proceeded with a repeated-measures ANOVA to evaluate differences in MEP amplitudes. The analysis revealed a significant effect of coil placement on MEP amplitudes (*F_(6, 54)_* = 7.75, *p* < 0.0001). Further post-hoc analyses using repeated t-tests with Holm-Bonferroni correction showed that MEP peak-to-peak amplitudes were significantly higher when coil placements were defined by the magnitude and neuron models (mean_mag_ = 0.737 mV, mean_neuron_ = 0.732 mV; *t_mag vs. neuron_* = 0.150, *p* = 0.884) compared to both the cosine model (mean_cos_ = 0.502 mV; *t_mag vs. cos_* = 4.016, *p* = 0.006; *t_neuron vs. cos_* = 4.545, *p* = 0.004), and the four adjacent control positions (mean_ant_ = 0.361 mV, mean_pos_ = 0.386 mV, mean_sup_ = 0.397 mV, mean_inf_ = 0.218 mV; *t_mag vs. controls_* = 4.445, *p* = 0.0032; *t_neuron vs. controls_* = 5.034, *p* = 0.0021). These findings suggest that the magnitude and neuron models more accurately identified the cortical areas responsible for generating MEPs in the target hand muscle, leading to more effective stimulation. Notable, four subjects even exhibited identical optimized coil positions for the magnitude and neuron models (Fig. 6B).

## 4. Discussion

TMS offers a non-invasive way to map motor functions in the brain. Early mapping studies relied on projecting coil positions with high MEP amplitudes directly onto the cerebral cortex (Wassermann et al., 1996; Classen et al., 1998; Krieg et al., 2014), but they were unable to identify target areas precisely. This is because simple projections fail to account for the complex shape of the brain, which affects the distribution of the induced E-field. Later studies using E-field simulations proposed that a cortical location could be classified as a certain muscle representation if the maximum E-field in that area evoked a response at the targeted muscle (Thielscher and Kammer, 2002). However, TMS reaches broader regions of the brain rather than a single activation point, leading to a less precise estimation of functionally relevant areas. A more promising direction involves correlating calculated E-fields with graded behavioral or physiological responses to better understand the mechanisms behind TMS effects.

Recent methods have improved TMS mapping by directly linking the induced E-field to observable MEP amplitudes in a sigmoidal pattern, resulting in more accurate and reliable cortical localizations. These studies have shown that the magnitude and normal component of the E-field (|𝐸| and |𝐸_⊥_|) might be the potential drivers of cortical stimulation (Weise et al., 2020; Numssen et al., 2021; Jing et al., 2023; Hikita et al., 2023). However, while including macroscopic properties like gyrification patterns, these localization approaches neglect the mesoscopic detail, such as the complex morphology of cortical neurons and their spatial organization within layers throughout the cortical gray matter. This neuronal architecture plays a crucial role in how the E-field interacts with neurons, influencing local cortical activation.

In this study, we extended the regression-based TMS mapping approach by incorporating an average response model that provides pre-calculated firing thresholds for various neuron types and orientations. By adjusting the |E| based on neuron-specific firing thresholds, a more refined understanding of how TMS affects neural circuits and influences plasticity processes should be provided.

### 4.1. Cortical Localization: Magnitude and Neuron Models Outperform the Cosine Model

We performed a sigmoidal regression analysis to localize the muscle representation of TMS-elicited MEPs. The comparison between the magnitude, neuron, and cosine models offers a deeper understanding of how different E-field quantities influence localization accuracy.

The ***magnitude*** and ***neuron models*** (incorporating neuron morphologies from cortical L2/3 and L5) demonstrated similar localization results, both in terms of the overall shape of the 𝑅^2^ maps, the 𝑅^2^peak values, and the locations of 𝑅^2^ peaks. This indicates that accounting for neuron-specific factors derived from an average response model does not significantly alter the overall localization compared to using the magnitude of the E-field alone. In other words, the magnitude model alone is already an effective, simple, and fast approach in identifying the relevant cortical sites responsible for the MEPs. Results showed that the FDI muscle representation is primarily located on the rim and the crown of the precentral gyrus, in accordance with previous studies (Bungert et al., 2017; Aonuma et al., 2018).

In contrast, the ***cosine model***, which assumes that stimulated axonal segments are exclusively oriented orthogonally to the cortical surface, explained significantly less variance in the regression compared to both the magnitude and neuron models. The consistently lower 𝑅^2^ values across all subjects indicate that the cosine model is less effective at capturing the functional stimulation site compared to the other two models. Moreover, 𝑅^2^ maps generated by the cosine model lacked the spatial coherence seen in the magnitude and neuron models, indicating more noise was introduced into the localization process. The distinct locations of maximum 𝑅^2^ values between cosine and the other two models further confirmed this. Accordingly, we do not recommend the cosine model to be used in future mapping studies.

In fact, the E-field strength is generally higher in the gyral crown, which is closer to the scalp, while the normal component of the E-field (as captured by the cosine model) is more prominent in the sulcal wall. This observation aligns with our results, showing distinct localization pattern differences between the cosine model and the other two models. Notably, functional MRI studies have found that TMS activates deeper cortical regions, such as the sulci, primarily due to proprioceptive sensory feedback from muscle twitches. When this feedback was suppressed in concurrent TMS-fMRI studies, however, the activation shifted closer to the gyral crown—areas of higher E-field magnitude rather than the normal component emphasized by the cosine model (Shitara et al., 2013). This supports our conclusion that compared to the cosine model, the magnitude model is better suited for mapping the cortical sites responsible for MEPs.

### 4.2. Convergence Analysis: Magnitude and Neuron Models Demonstrate Faster Convergence

The convergence analysis revealed significant differences in how quickly each model achieves stable cortical localization with respect to the number of TMS trials. Both the magnitude and neuron models demonstrated faster convergence compared to the cosine model, as indicated by the geodesic distance of 𝑅^2^ peak locations and the overall similarity 𝑅^2^maps measured by NRMSD.

In terms of the geodesic distance compared to the ground truth (N trials), which evaluates the precision of peak localization, the magnitude and neuron models required far fewer trials (309 and 282, respectively) to reach a 1 mm threshold compared to the cosine model, which needed 622 trials. However, no significant difference was found between the magnitude and neuron models, suggesting that neuron-specific adjustments do not drastically improve convergence speed over the magnitude model alone. Similarly, the NRMSD analysis, which assesses the overall similarity between 𝑅^2^ maps, further supported the superiority of the magnitude and neuron models. Both required fewer than 200 trials to reach a 5% deviation threshold, while the cosine model needed 278 trials to achieve the same stability.

In summary, these results emphasize that both the magnitude and neuron models outperform the cosine model in terms of convergence speed and efficiency, making them better choices for future TMS studies. Given they converged at similar rates, the magnitude model stands out as a simpler and faster approach, offering efficient localization without the need for complex neuron-specific adjustments. We recommend at least 300 TMS trials are needed for the regression-based TMS mapping to get stable localization results.

### 4.3. Validation: Magnitude and Neuron Models Produce Higher MEP Amplitudes

The validation experiment in our study aimed to determine which model most accurately locates the cortical hotspots responsible for eliciting MEPS in the FDI hand muscle. The results demonstrated that both the magnitude and neuron models produced significantly higher peak-to-peak MEP amplitudes compared to the cosine model and the adjacent control positions. Moreover, 4 subjects even exhibited identical optimized coil positions for the magnitude and neuron models, further reinforcing their similarities in cortical localization. These findings validate the superiority of the magnitude and neuron models over the cosine model in pinpointing the cortical regions involved in generating MEPs. The lower MEP amplitudes observed in the cosine model and control positions suggest that these locations are less functionally relevant for eliciting strong MEP responses.

## 5. Conclusions

In this study, we applied a regression-based localization method to identify the cortical representation of muscle responses to TMS. We focused on modeling the induced E-field across different coil placements to investigate whether its distribution could explain physiological observations related to MEPs. We evaluated three different models for cortical localization: the magnitude model (regardless of E-field direction), the neuron model (accounting for cortical neuron morphology), and the cosine model (overemphasizing the directional sensitivity of neurons). This is the first study to attempt cortical mapping that integrates a realistic model of cortical neuron morphology and systematically compares results with traditional neuronal response models.

Our findings demonstrated that on the motor cortex, both the magnitude and neuron models consistently outperformed the cosine model in terms of precision, efficiency, and stability. Their faster convergence and higher accuracy in identifying cortical hotspots highlight their practical advantages for TMS studies. These findings imply that the simpler magnitude model is already highly effective and offers the added benefit of reduced computational demands, making it an accessible and robust choice for TMS mapping. The neuron model, however, represents a promising approach for integrating cortical neuron morphology, and may potentially enhance cortical mapping directly with respect to simulated cortical response functions, so long as detailed cell-layer data is available. When the cytoarchitectural details of a target region are unknown, our study suggests that the magnitude model could serve as a reasonable first approximation.

We, therefore, recommend the simpler magnitude model for future TMS mapping research in the motor cortex, requiring at least 300 trials for stable and reliable localization results. The cosine model, however, due to its slower convergence and lower precision, is not recommended for future studies.

